# Phenotypic Switching of Vascular Smooth Muscle Cells in Duchenne Muscular Dystrophy

**DOI:** 10.1101/2023.06.23.546309

**Authors:** Wanling Xuan, Feng Cheng, Xiaowei Han, Srinivas M. Tipparaju, Muhammad Ashraf

## Abstract

**Background:** Extensive studies have been conducted in skeletal muscle and myocardium affected by Duchenne Muscular Dystrophy (DMD) disease but there is a significant gap of research in the role of vascular smooth muscle cells (VSMCs) in DMD. Here, we investigated the role of dystrophin deficiency in the maintenance of VSMCs contractile phenotype.

**Methods:** 12-14 months old mdx mice and DMD induced pluripotent stem cells (iPSC) derived VSMCs were used as disease models. Morphological and immunohistochemistry analyses were performed to determine histological changes and the expression of contractile markers. Transmission Electron Microscopy (TEM) was used to assess ultrastructural changes in the VSMCs. Mito-tracker staining and TUNEL staining were performed to determine mitochondria fission-fusion and apoptosis respectively. mRNA Sequencing for normal iPSC derived VSMCs (WT-VSMCs) and DMD iPSC derived VSMCs (DMD-VSMCs) with or without oxidative stress was performed. KEGG signaling pathway enrichment, Go function enrichment and Gene set enrichment analysis (GESA) were conducted to explore the potential mechanism responsible for these changes. In addition, transcription factor enrichment analysis was performed to unravel mechanistic pathways of regulatory networks.

**Results:** Spontaneous abnormal VSMCs proliferation, loss of vascular structure and degenerative changes occurred in VSMCs in aorta from 12-14 months old mdx mice. The DMD-VSMCs showed maturation defect, loss of mitochondrial hemostasis, and increased vulnerability to oxidative stress compared with WT-VSMCs. Transcriptome analysis revealed dysregulation of smooth muscle proliferation, differentiation, and vascular development in DMD-VSMCs. Transcriptional factor, target, and motif discovery analysis of the dysregulated gene set suggested potential contributions of transcriptional factors GADD45A, SOX9, TIA1, RBBP9 and FOXM to the phenotypes of DMD-VSMCs. Under oxidative stress, initiation of apoptotic process was significantly enhanced in DMD-VSMCs while their response to hypoxia and oxidative stress was downregulated.

**Conclusions:** Dystrophin deficiency induced VSMCs phenotype switching and disrupted mitochondrial metabolism. The findings in this study underscore the importance of vascular dysfunction in DMD disease and therapeutic interventions to restore VSMC phenotype may ameliorate the propensity of disease progression. It is suggested that the transcriptome analysis may allow the discovery of potential signaling pathways involved in the dysregulation of transcription factors.

## Introduction

Dystrophin is critical for maintaining the integrity of muscle cell membranes ^1, 2^. A previous clinical study demonstrated that blood loss in patients with Duchenne Muscular Dystrophy (DMD) was much higher than that in patients with spinal muscular atrophy undergoing same or similar surgery, supporting the possibility that a poor vascular smooth muscle cells (VSMCs) vasocontractile response may be due to a lack of dystrophin ^3^. Furthermore, reduction of intramuscular blood flow has been observed in DMD patients, suggesting an existence of functional muscle ischemia ^4^. Potential vascular dysfunction has been reported in DMD disease ^5-7^. This vascular defect was considerably reduced in DMD mouse by improving angiogenesis with overexpression of growth factors ^8^ and reduction of intramuscular blood flow has been reported in DMD patients ^4^. Recently, it has been reported that both vasoconstriction and vasorelaxation were comprised in dystrophin deficient dogs ^9^. Vascular dysfunction blunts the blood supply, impairs muscle tissue perfusion, causes muscle necrosis and degeneration, and limits gene therapy delivery. Thus, vascular defects may be an underlying pathogenic mechanism in DMD disease. However, the role of dystrophin deficiency in VSMCs has not been thoroughly studied.

The most common DMD mouse model is mdx mice, which carry a nonsense mutation in exon 23 of the dystrophin gene ^10,11^. The association between muscle structure and function with age throughout the life span of the mdx mice has only recently been appreciated ^12^. However, despite the absence of dystrophin in muscles, adult mdx mice do not exhibit the pathogenic progression characteristic of human DMD, severe muscle weakness, loss of muscle weight, accumulation of fat and fibrosis is not significant until almost two years of age ^11^. Furthermore, an age-dependent effect on angiogenesis in mdx mice has been reported ^13^. Significant vessel density is decreased in skeletal muscle from 12-month-old mdx mice compared with 3-month-old mdx mice and aged match wild type mice ^5^. A previous study reported that isometric force was decreased in VSMCs from mdx mice after nitric oxide stimulation ^14^. However, the authors did not reveal the specific age of the mdx mice they used in the study. Upon mechanical injury, increased neointima formation and smooth muscle proliferation were observed in 4-5 months old mdx mice. However, they did not report spontaneous neointima thickness in these mdx mice. As previously reported that significant vessel defects were observed in 12-month-old mdx mice, here we investigated whether any spontaneous phenotypic changes take place in VSMCs from 12-14 months old mdx mice. Decreased dystrophin expression was reported in synthetic smooth muscle cells compared with contractile smooth muscle cells in vitro ^15^. Similarly, stabilization of actin filaments promoted the expression of dystrophin ^14^ in mouse aorta smooth muscle cells. However, whether dystrophin deficiency promotes phenotypic switching, and the underlying mechanism remains unknown. Human induced pluripotent stem cells (iPSC)-based disease models are promising due to the availability of unlimited supply of clinically relevant phenotypic cells of human origin in rare diseases such as DMD. Thus, in this study, we utilized DMD patient derived iPSC to differentiate into VSMCs to study their phenotype changes in comparison with normal iPSC derived VSMCs.

Here, we found spontaneous abnormal proliferation of VSMCs, loss of vascular structure and degenerative changes of VSMCs in aorta from 12-14 months old mdx mice. The DMD iPSC derived VSMCs (DMD-VSMCs) showed maturation defects and increased vulnerability to oxidative stress compared with wild type VSMC. Transcriptome analysis revealed dysregulation of smooth muscle proliferation and differentiation and vascular development related biological function in DMD-VSMCs. Under oxidative stress, apoptotic process was significantly enhanced in DMD-VSMCs while their response to hypoxia and oxidative stress was downregulated. Transcriptional factor, target, and motif discovery analysis of the dysregulated gene set suggested potential contributions of transcriptional factors GADD45A, SOX9, TIA1, RBBP9 and FOXM to the phenotypes of DMD-VSMCs.

## Methods

### Human iPSC culture

The Human iPSC cell lines DYS0100 from ATCC Company and human DMD iPS cell line SC604A from SBI Company were used. iPSC were cultured on vitronectin coated six-well plate in mTeSR1 medium (Stem Cell Technologies) with a daily change. iPSC were passaged using ReLeSR™ passaging reagent (Stem Cell Technologies).

### Generation of vascular smooth muscle cells from iPS cells and their characterization

Human iPSC at passages 20–30 were used for vascular smooth muscle cells (VSMCs) differentiation. The differentiation protocol outline is shown in **Fig.1A**. Briefly, human iPSC were cultured on vitronectin using mTeSR1 medium. Upon confluency, the medium was switched to α-MEM basal medium supplemented with 20% Knock out serum (KSR), 1mM L-Glutamine, 10mM Nonessential Amino Acid and 10μM SBB-431542 for 10 days. Next, the cells were trypsinized and seeded at a density of 4×10^4^ cells/cm^2^ onto uncoated culture dishes in expansion medium (α-MEM basal medium+10% FBS). After the third passage, a morphologically homogeneous population of mesenchymal stem cells (MSC)-like cells became evident and were tested for mesenchymal makers. Then the MSC-like cells were further differentiated into VSMCs using 5ng/ml TGFβ treatment for 6 days. The differentiated VSMCs were characterized with α-SMA, calponin, SM-22α and Myh11 staining. For immunofluorescence staining, cells were fixed with 4% PFA for 10 mins and blocked with 10% FBS for 1h at room temperature. Cells were incubated with primary antibodies including anti-CD105 (sc-18838, Santa Cruz), NG2 (MAB5384A4, Millipore Sigma), α-SMA (ab5694, Abcam), calponin (C2687, Sigma), SM- 22α (10493-1-AP, Thermo Fisher Scientific) and Myh11(NBP1-87025, Novus Biologicals) respectively at 4 °C overnight and secondary antibody conjugated to Alexa Fluor 594 or Alexa Fluor 488 (Life Technologies) at room temperature for 1 h. DAPI was used as nuclear counterstain.

**Fig. 1.**
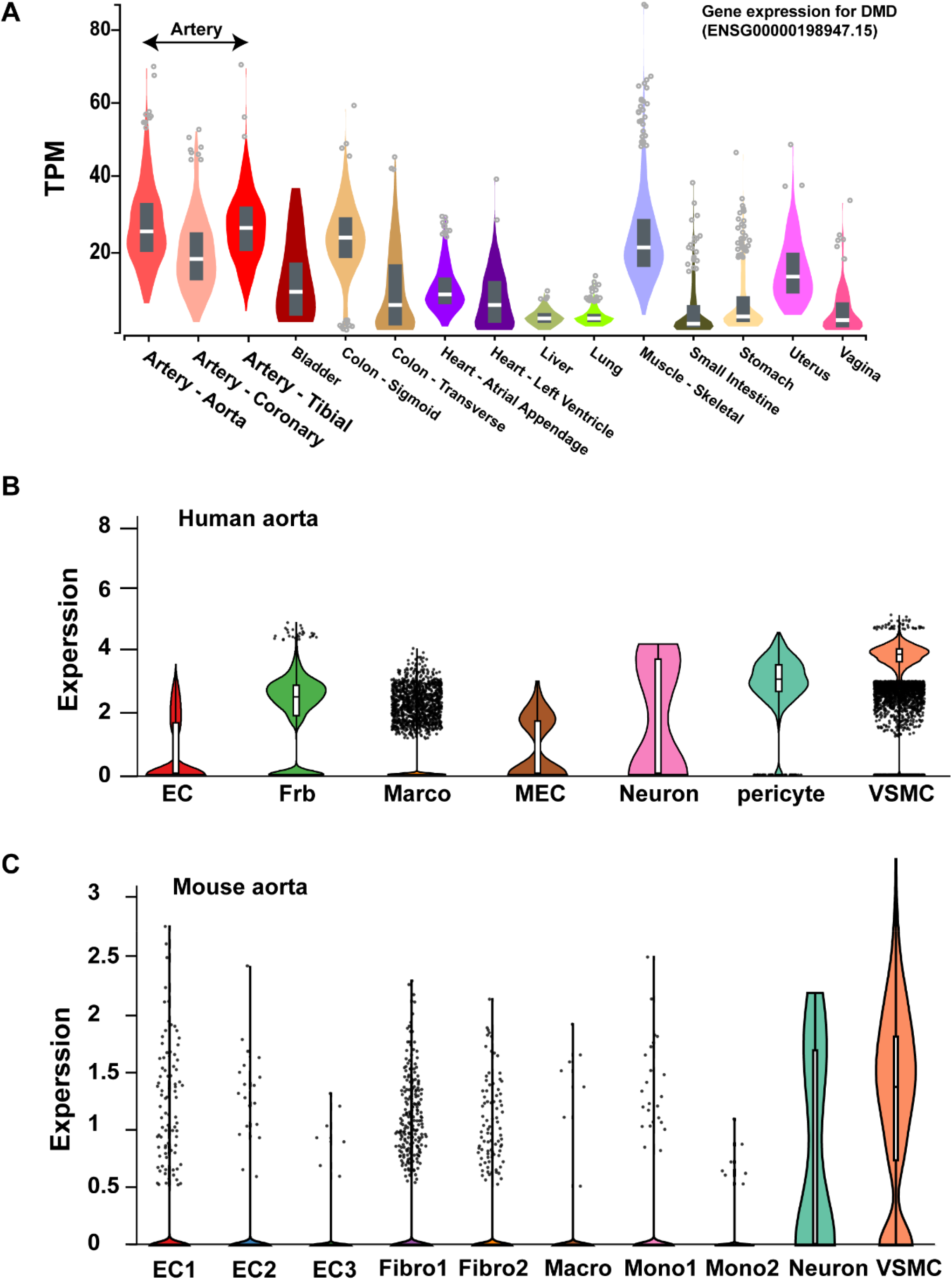
DMD gene expression in different tissue and cell types. (A) DMD gene expression in different tissues. High expression of DMD gene in arteries. Data from GTx portal database. High expression of DMD gene in VSMCs from human aorta (B) and mouse aorta (C) revealed by scRNA-seq (Broad Institute Single Cell Portal). EC: endothelial cell; Frb/Frbio: fibroblast; Mono: monocyte; Macro: macrophage; VSMC: vascular smooth muscle cell.

### Oxidative stress

VSMCs differentiated from normal iPSC or DMD iPSC were subjected to 100 μM hydrogen peroxide (H_2_O_2_) treatment for 24h for oxidative stress.

### Histology

Mouse aortic tissue from 12-14 months old wild type mice (Stock No: 000665 The Jackson Laboratory) and mdx mice (Stock No: 018018, The Jackson Laboratory) was harvested and processed for Masson’s trichrome staining as reported previously ^16^. Animal experiments were carried out according to the experimental protocol approved by the Augusta University Animal Care and Use Committee (approval number 2018-0940).

### Immunohistochemistry analysis

Mouse aortic tissue was collected and fixed with 4% paraformaldehyde (PFA) for 1 h at room temperature and then immersed in 30% sucrose overnight at 4 °C as described in our previous study ^16^. On second day, the aortic tissue was embedded in an optical cutting temperature (OCT) compound (Tissue Tek), and then the samples were sliced into 5-μm-thick frozen cross-sections using a Leica CM3050 cryostat. Sections were incubated with anti-Myh11 (NBP1-87025, Novus Biologicals) at 4 °C overnight respectively and anti- mouse secondary antibody conjugated to Alexa Fluor 488 (Life Technologies) at room temperature for 1 h. Images were acquired with an Olympus confocal microscope.

### Transmission Electron Microscopy

Mouse aorta was processed for electron microscopy to assess ultrastructural changes in the VSMCs from mdx mice. Samples were processed and imaging was carried out by the Augusta University histology and TEM core. Briefly, aortic tissues were cut into 1mm^3^ pieces and fixed in 2.5% glutaraldehyde in 0.1 M sodium cacodylate buffer (pH 7.4). Next, tissues were post-fixed in 1% osmium tetroxide in 1% K4Fe (CN)6 buffer with 0.1 M sodium cacodylate, dehydrated through a graded series of ethanol and propylene oxide, and embedded in Epon 812. Ultrathin cryo-sections were prepared using Leica UC7 Ultramicrotomes, mounted on copper grids and stained with lead citrate and uranyl acetate. Images were captured by JEOL JEM-1230 Transmission Electron Microscope.

### Mito-tracker staining

Mitochondria in cultured cells were visualized by staining with MitoTracker Red CMXRos dye (Thermo Fisher Scientific). The dye was added to the live cells at a final concentration of 100 nM and incubated at 37°C for 30 minutes. Cells were washed and fixed in 2% PFA in PBS. Images were taken by a confocal microscope (Olympus, Japan).

### TUNEL staining

Terminal deoxynucleotidyl transferase-mediated dUTP nick-end labeling (TUNEL) was performed to assess cell apoptosis using commercial apoptosis detection kit (Thermo Fisher Scientific, United States). All procedures were done according to the directions of the manufacturers. Cells were counterstained with DAPI to visualize nuclei. The number of TUNEL-positive cells was determined by randomly counting 10 fields and was expressed as a percentage of the cells with normal nuclei.

### Western Blot

Cell extracts were lysed with radio immunoprecipitation assay (RIPA) buffer supplemented with Protease Inhibitors Mixture (Sigma). Pierce™ BCA Protein Assay Kit (Thermo Scientific) was used to determine protein concentration. 10μg protein was separated by SDS/PAGE and transferred to the PVDF membrane (Bio-Rad). Membranes were incubated with primary antibodies overnight at 4°C: α-SMA (ab5694, abcam), Calponin (C2687, Sigma) and Tubulin (2128S, Cell Signaling Technology). On second day, membranes were incubated with an anti-mouse/rabbit peroxidase-conjugated secondary antibody at room temperature. Immunoreactive bands were visualized by the enhanced chemiluminescence method (Bio-Rad) with Fluorchem E detection system (ProteinSimple USA). The relative expression levels of target proteins were quantified by ImageJ software (National Institutes of Health, Bethesda, MD, United States).

### mRNA Sequencing and Analysis

mRNA sequencing was performed by genomic core of Washington University in St. Louis. Samples were prepared according to library kit manufacturer’s protocol, indexed, pooled, and sequenced on an Illumina NovaSeq 6000. Basecalls and demultiplexing were performed with Illumina’s bcl2fastq software and a custom python demultiplexing program with a maximum of one mismatch in the indexing read. RNA-seq reads were then aligned to the Ensembl release 76 primary assembly with STAR version 2.5.1a1. Gene counts were derived from the number of uniquely aligned unambiguous reads by Subread: featureCount version 1.4.6-p52. Isoform expression of known Ensembl transcripts were estimated with Salmon version 0.8.23. Sequencing performance was assessed for the total number of aligned reads, total number of uniquely aligned reads, and features detected. The ribosomal fraction, known junction saturation, and read distribution over known gene models were quantified with RSeQC version 2.6.24. All gene counts were then imported into the R/Bioconductor package EdgeR5 and TMM normalization size factors were calculated to adjust for samples for differences in library size. Ribosomal genes and genes not expressed in the smallest group size minus one sample greater than one count-per-million were excluded from further analysis. The TMM size factors and the matrix of counts were then imported into the R/Bioconductor package Limma6. Weighted likelihoods based on the observed mean-variance relationship of every gene and sample were then calculated for all samples with the voomWithQualityWeights7.The performance of all genes was assessed with plots of the residual standard deviation of every gene to their average log-count with a robustly fitted trend line of the residuals. Differential expression analysis was then performed to analyze differences between conditions and the results were filtered for only those genes with Benjamini-Hochberg false-discovery rate adjusted *p*-values less than or equal to 0.05.

For the different expression genes (DEGs) with 2-fold changes, KEGG pathway enrichment was performed using DAVID online tools.; Go categories of biological process (BP), molecular function (MF) and cellular components (CC) were performed using the online Go enrichment analysis tools (http://geneontology.org). The upregulated DEGs and downregulated DEGs were entered separately. *P<*0.01 and FDR <0.05 were as the cutoff criteria to identify the enriched pathways. Results were plotted by https://www.bioinformatics.com.cn (last accessed on 20 Feb 2023), an online platform for data analysis and visualization. Gene set enrichment analysis (GSEA) tools (java GSEA Desktop Application version 4.3.2; http://software.broadinstitute.org/gsea/downloads.jsp) were also used to perform enrichment analysis of KEGG pathway and Go categories for DEGs. The significantly enriched pathways and Go terms were defined by nominal *P<0*.*01* and FDR<0.25. Transcriptional factor, target, and motif discovery analysis of upregulated genes with 2-fold changes was performed using iRegulon plugin in Cytoscape_3.9.1 software.

DMD expression in different tissue was analyzed from data in GTx portal database. DMD gene expression in VSMCs form human aorta dataset (https://singlecell.broadinstitute.org/single_cell) and mouse aorta dataset (https://singlecell.broadinstitute.org/single_cell/study/SCP1361/single-cell-transcriptome-analysis-reveals-cellular-heterogeneity-in-the-ascending-aorta-of-normal-and-high-fat-diet-mice) was analyzed using Broad Institute Single Cell Portal.

### Statistical analysis

Data were expressed as mean ± SD. Normality was tested and statistical analysis of differences among the different groups was compared by unpaired two-tailed Student’s t-tests. The differences were considered statistically significant at *P < 0*.*05*. Statistical analyses were performed by Graphpad Prism 9.5.

### Results

### DMD expression in VSMCs

Tissue expression analysis showed an abundant expression of dystrophin in arteries from GTEx portal database (**Fig.1A**). Analysis of single cell RNA-seq data of normal adult mouse thoracic aorta and human aorta, dystrophin was highly expressed in the VSMCs compared with other cell types in the aorta (**Fig.1B and 1C**).

### Phenotypic changes in VSMCs from mdx mice

Gross images of aorta from wild type and mdx mice are shown. Histology analysis and TEM images showed an abnormal proliferation of VSMCs, loss of vascular structure and degenerative changes of VSMCs (Intranuclear and cytoplasmic vacuoles) in aorta from 12-14 months old mdx mice (**Fig.2**). A significant decrease of myosin heavy chain (Myh11) expression in aorta was also observed (**Fig.2**). These observations supported that dystrophin deficiency caused loss of contractile phenotype of VSMCs.

**Fig. 2.**
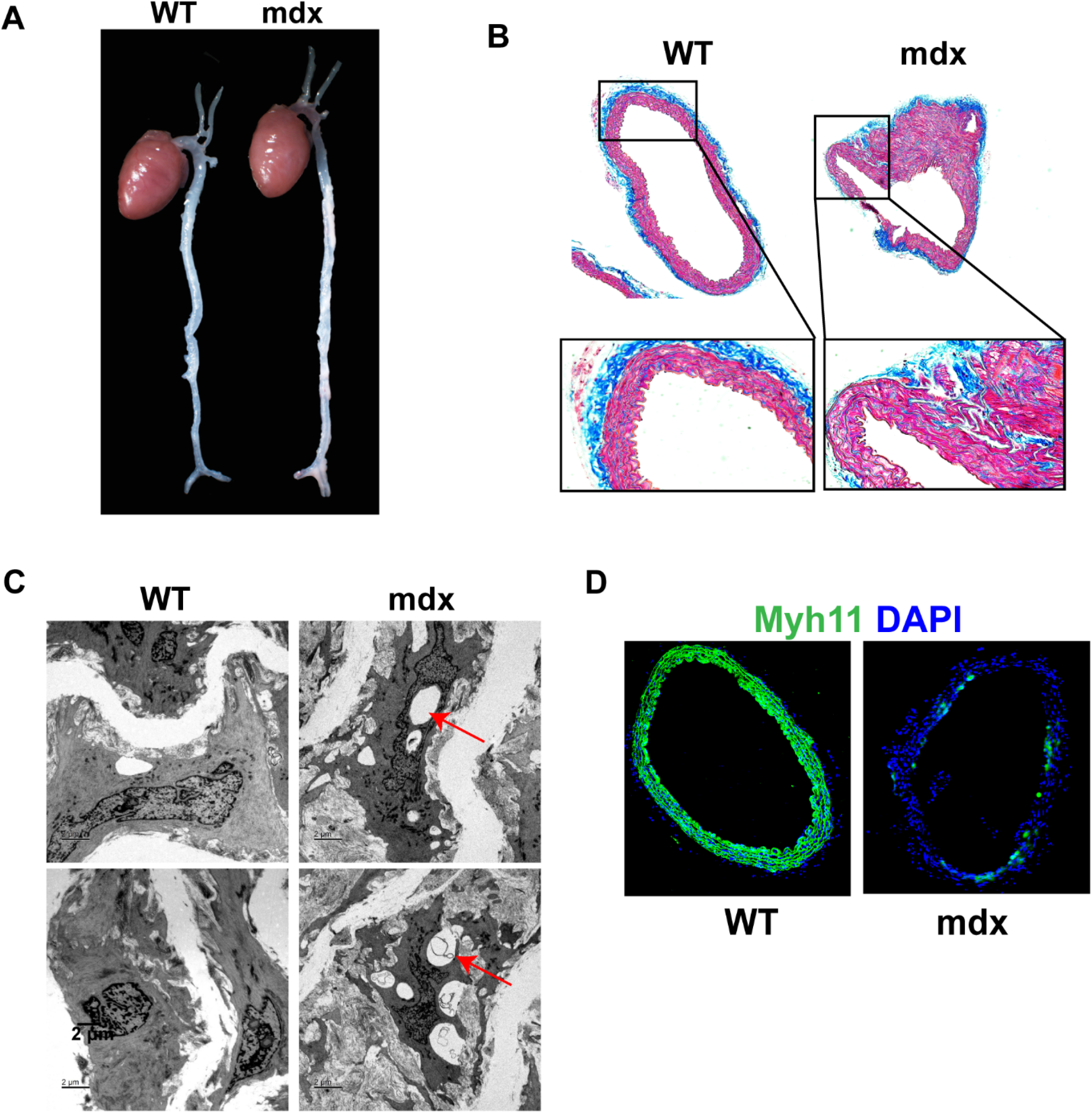
Phenotypic changes in VSMCs from mdx mice. (A) Gross morphology of heart and aorta from 12-14 months old wild type (WT) and mdx mice. (B) Increased proliferation of VSMCs and loss of vascular structure in aorta from 12-14 months old mdx mice shown by Trichome Masson staining. (C) TEM images showed degenerative changes of VSMCs (Intranuclear and cytoplasmic vacuoles, shown by arrows) in aorta from 12- 14 months old mdx mice. (D) Decreased contractile protein-Myh11 expression in aorta from 12-14 months old mdx mice compared with age matched WT mice. n=5.

### Characterization of vascular smooth muscle cells (VSMCs) from normal WT iPSC and DMD iPSC

Human iPSC-based disease models are promising due to unlimited supply of clinically relevant phenotypic cells of human origin, especially for rare diseases such as DMD. Accordingly, we generated VSMCs from human normal iPSC and DMD-iPSC (SC604A, MD) cell lines by treatment with a small molecule, SB43152 followed by transforming growth factor-β (TGF-β) (**Fig.3A**). With SB43152 treatment, iPSC were differentiated into mesenchymal stem cell (MSC)-like cells expressing MSC markers (NG2 and CD105) with negligible expression of α-SMA and negative calponin expression (**Fig.3B**). We further differentiated these cells into VSMCs using TGF-β1 as previously described ^17^ . The differentiated VSMCs expressed VSMCs markers including α-SMA, calponin and SM-22α (**Fig.3C**). However, compared with normal VSMCs, DMD iPSC derived VSMCs displayed non-mature phenotype with low expression of contractile proteins (Calponin, SM-22α and Myh11) either by immunofluorescence (**Fig.3C**) or Western blot (**Fig.3D**). Enhanced mitochondrial fission was observed in DMD-iPS cells derived VSMCs (**Fig.4**) and in VSMCs from 12 months old mdx mice (**Fig.4**). These results suggested that dystrophin deficiency caused loss of mitochondrial homeostasis in VSMCs. In addition, DMD deficiency VSMCs were vulnerable to oxidative stress with increased cell apoptosis compared with normal VSMCs (**Fig.5**).

**Fig. 3.**
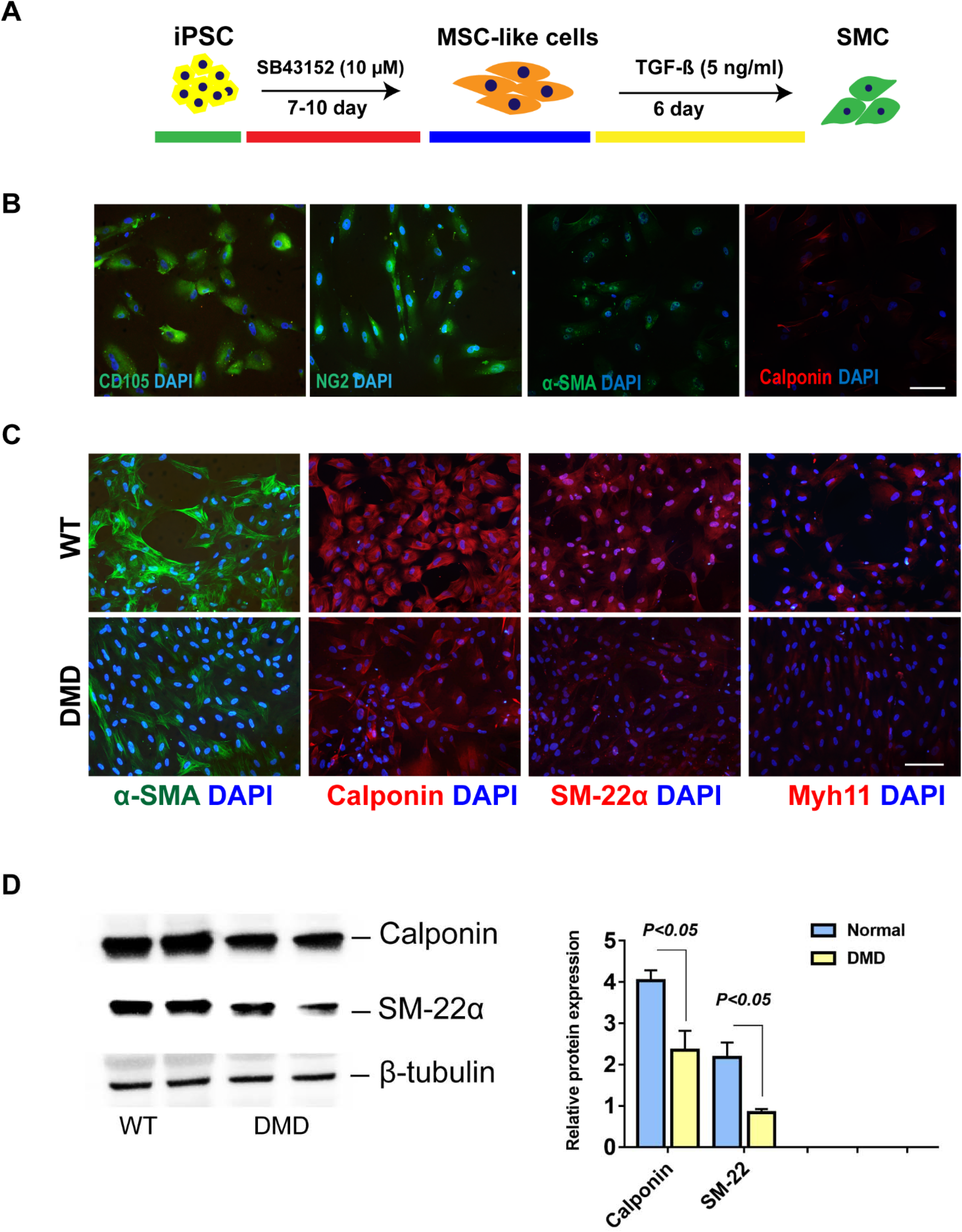
Generation of VSMCs from normal and DMD iPS cells. (A) Schematic outline of vascular smooth muscle cells (VSMCs) differentiation from iPSC; (B) Differentiated MSC-like cells expressed MSC markers: CD105 and NG2, with low expression of α-SMA, calponin expression is undetectable. (C) VSMCs markers expression by immunostaining in VSMCs derived from normal and DMD iPSC. (D) Representative Western blot images and semi-quantitative estimate of VSMCs markers expression. iPSC: Induced Pluripotent Stem Cells; MSC: Mesenchymal stem cells. n=3.

**Fig. 4.**
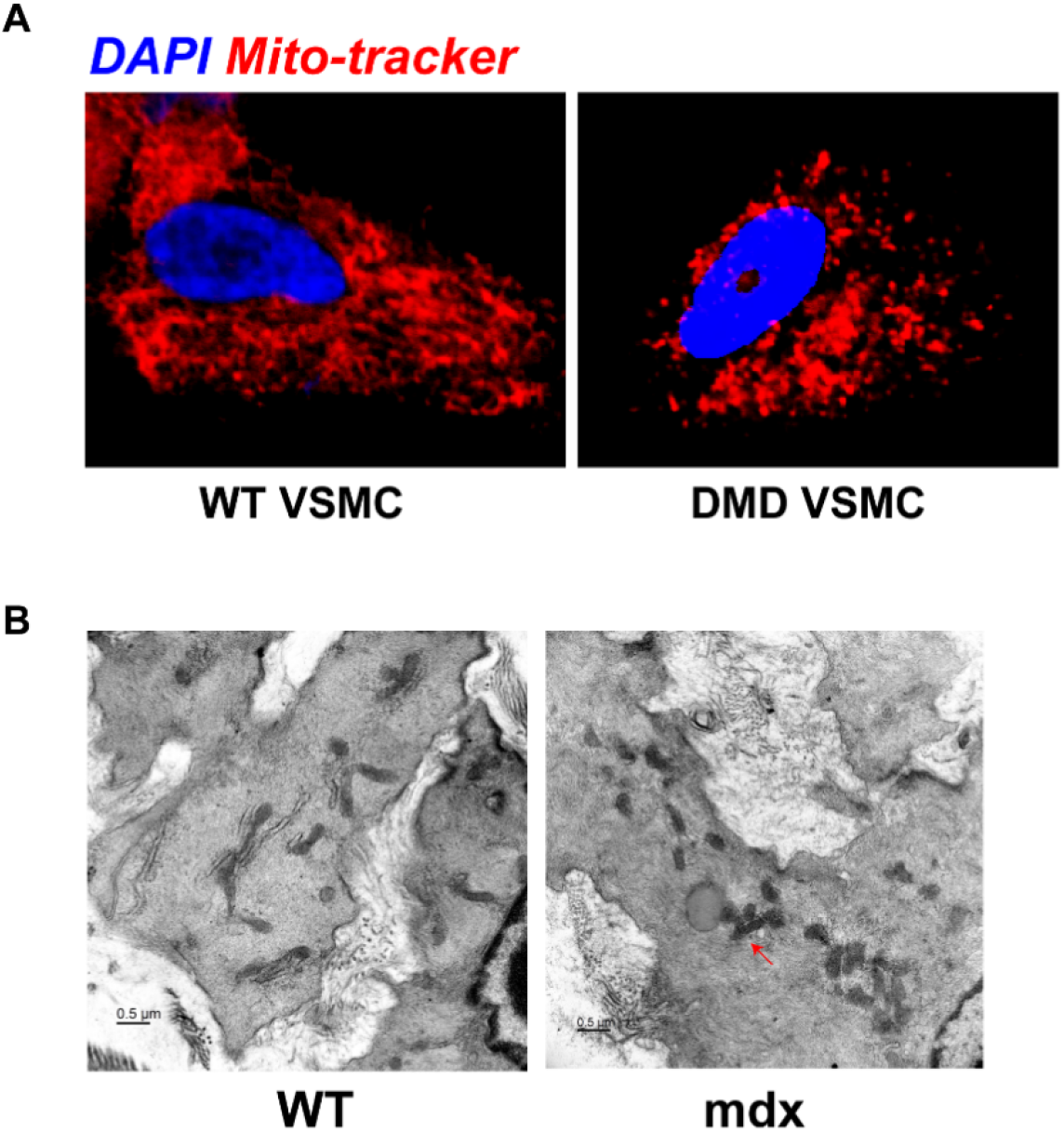
Abnormal mitochondrial fission in dystrophin deficiency VSMCs. (A) Enhanced mitochondrial fission in VSMCs derived from DMD PSC compared to wild type (WT) VSMCs. Mitochondria were identified by Mito-tracker. (B) TEM images showing mitochondrial fission in aortic VSMCs from 12 months old mdx mice. Mitochondria were small and fragmented (arrows). n=3.

**Fig. 5.**
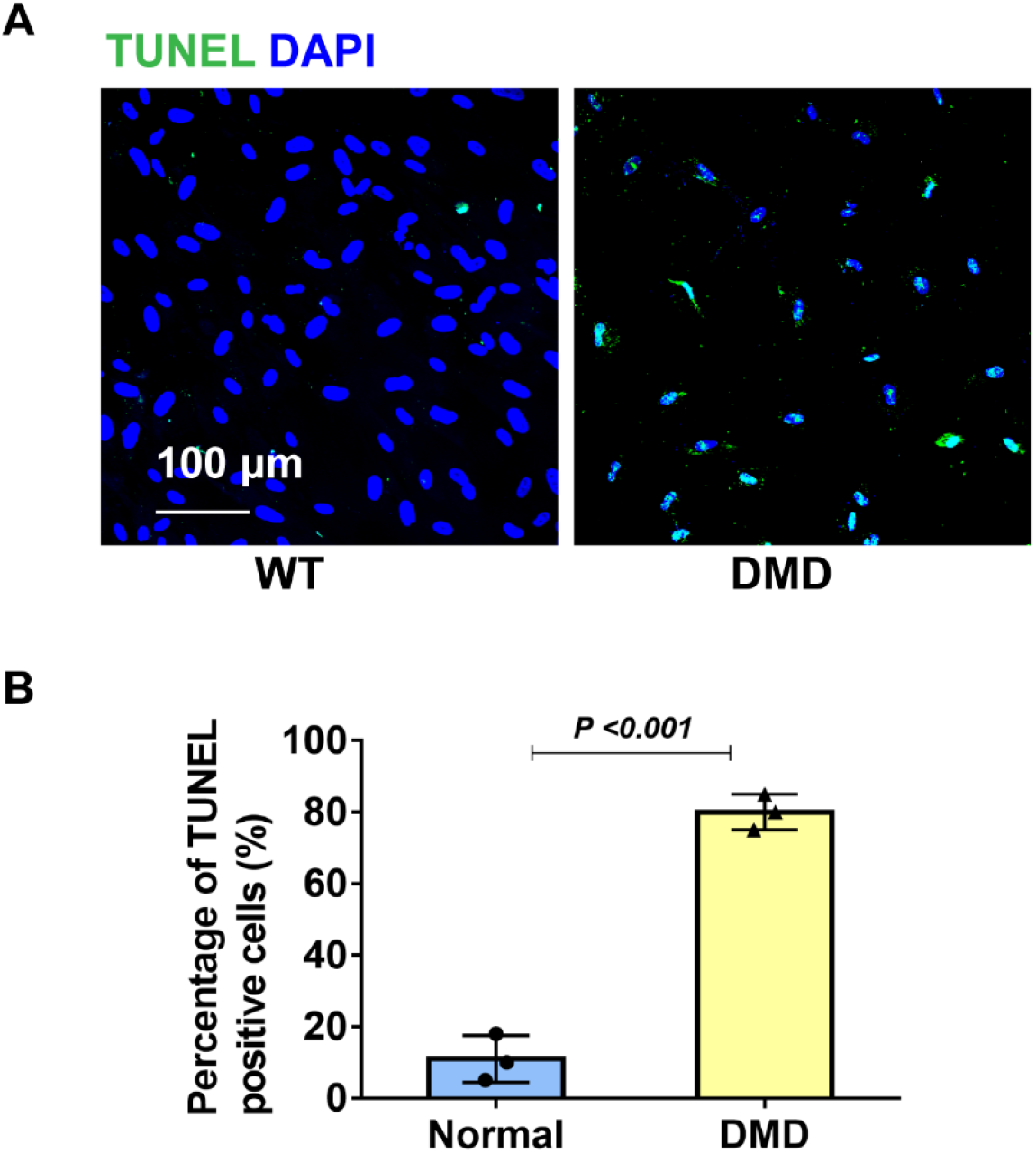
VSMCs from DMD iPSC is vulnerable to oxidative stress. (A) TUNEL positive cells in VSMCs from normal and DMD iPS cells after 24h H_2_O_2_ (100 μM) treatment. (B) Semi-quantitative estimate of TUNEL positive cells in VSMCs with H_2_O_2_ treatment. n=3.

### Transcriptome analysis of VSMCs from normal WT iPSC and DMD iPSC

Using DAVID tools, we performed KEGG enrichment analysis for VSMCs transcriptomic profiling of normal and DMD-iPSC derived VSMCs. DEGs were mainly enriched in upregulation of TGF-β, P53, Rap1 and Hippo signaling pathways and downregulation of ferroptosis in DMD-iPSC derived VSMCs (**Fig.6**). Go enrichment analysis showed negative regulation of smooth muscle cell differentiation and positive regulation of osteoblast differentiation in DMD-VSMCs (**Fig.7A**). Vascular development was also significantly decreased in dystrophin deficient VSMCs (**Fig.7B**). GSEA revelated enrichment of genes related to mitochondrial depolarization, and response to irons (**Fig.8**). Transcriptional factor, target, and motif discovery analysis of the dysregulated gene set showed enrichment of transcriptional factors (TFs) GADD45A, SOX9, TIA1, RBBP9 and FOXM1 (**Fig.9**).

**Fig. 6.**
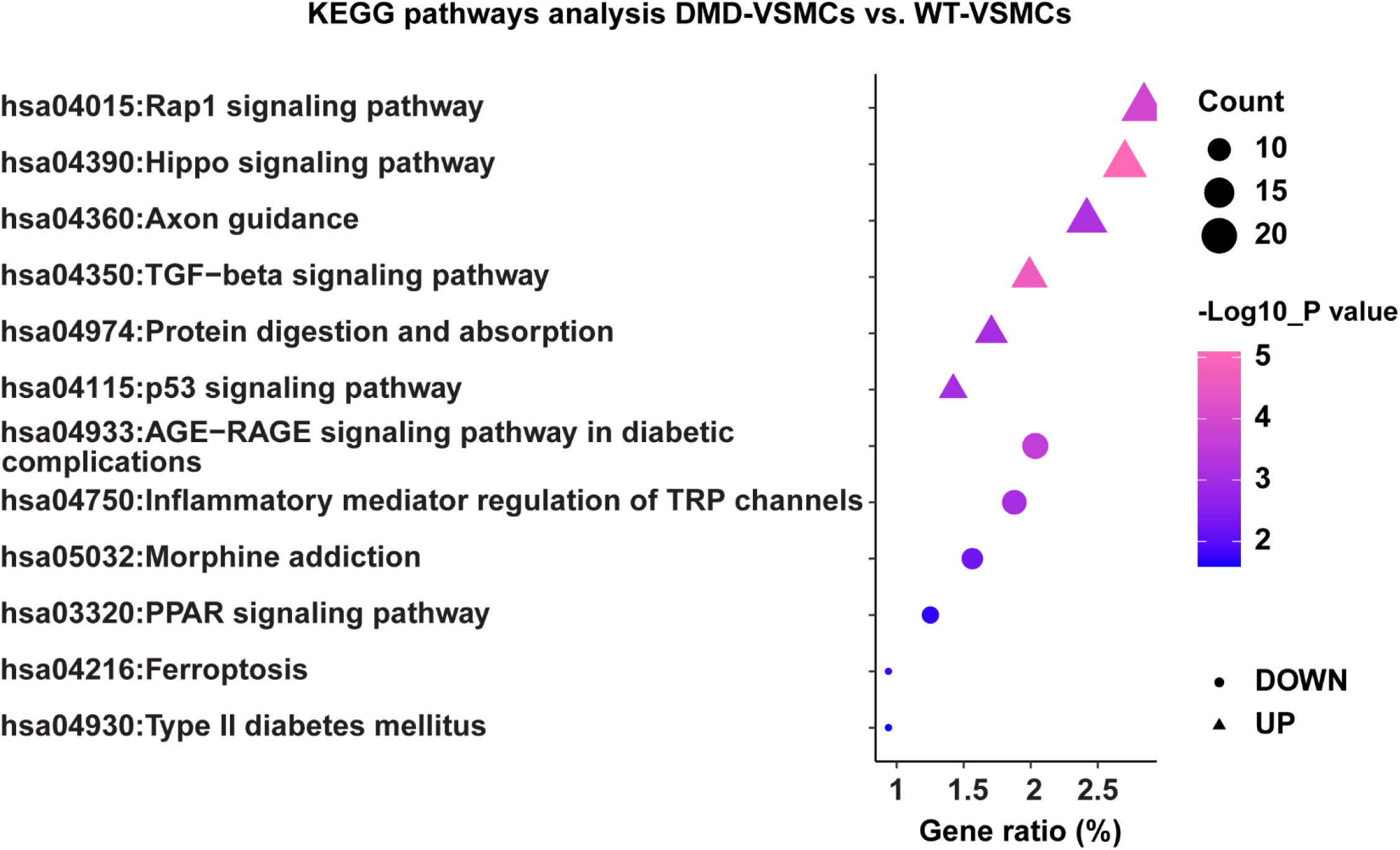
KEGG pathway analysis of fold enrichment from up and down regulated mRNAs with top 5 in DMD-VSMCs compared to WT-VSMCs.

**Fig. 7.**
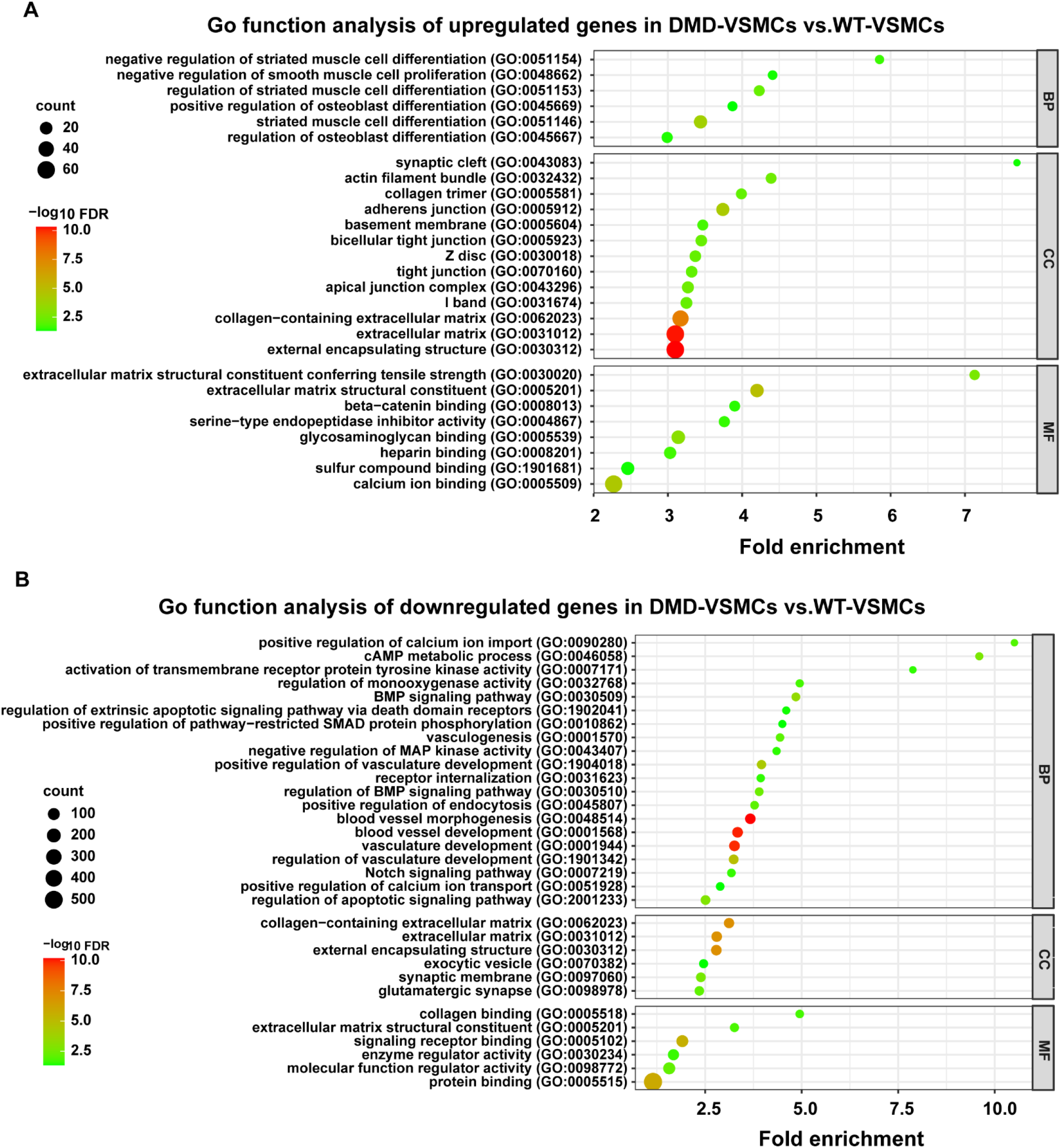
Go enrichment analysis for normal WT-VSMCs and DMD-VSMCs. Go categories of biological process (BP), molecular function (MF) and cellular components (CC) showing significant enrichment in the dataset of genes upregulated (A) and downregulated (B) in DMD-VSMCs compared to normal WT-VSMCs. Low -log_10_ (FDR) values are in green and high -log10 (FDR) values are in red, the size of the circle is proportional to the number of enriched genes.

**Fig. 8.**
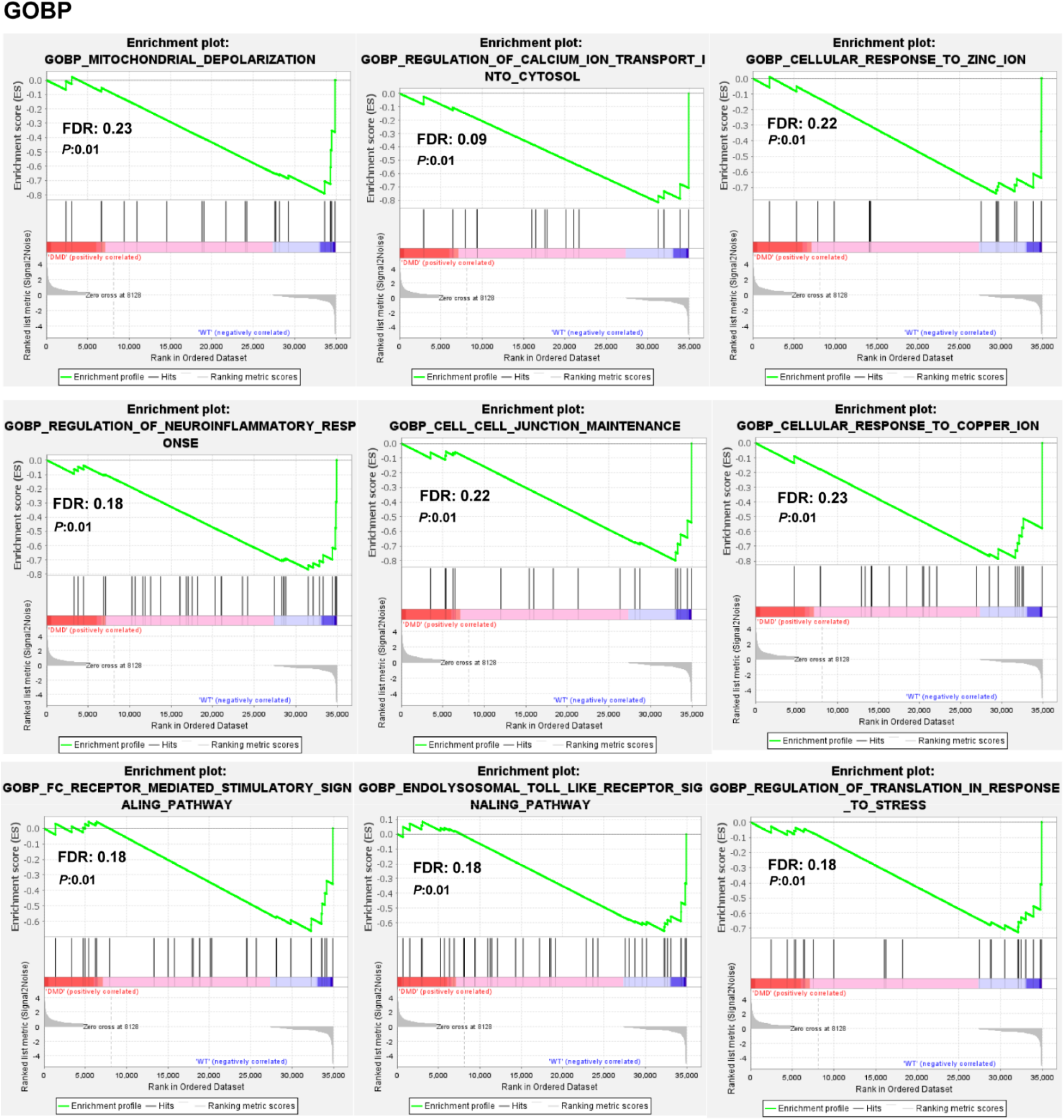
Gene set enrichment analysis (GSEA) for normal WT-VSMCs and DMD-VSMCs. GSEA biological process (BP) enrichment analysis showing significant enrichment in the dataset of genes downregulated in DMD-VSMCs (*P<0*.*01*, FDR≤0.25).

**Fig. 9.**
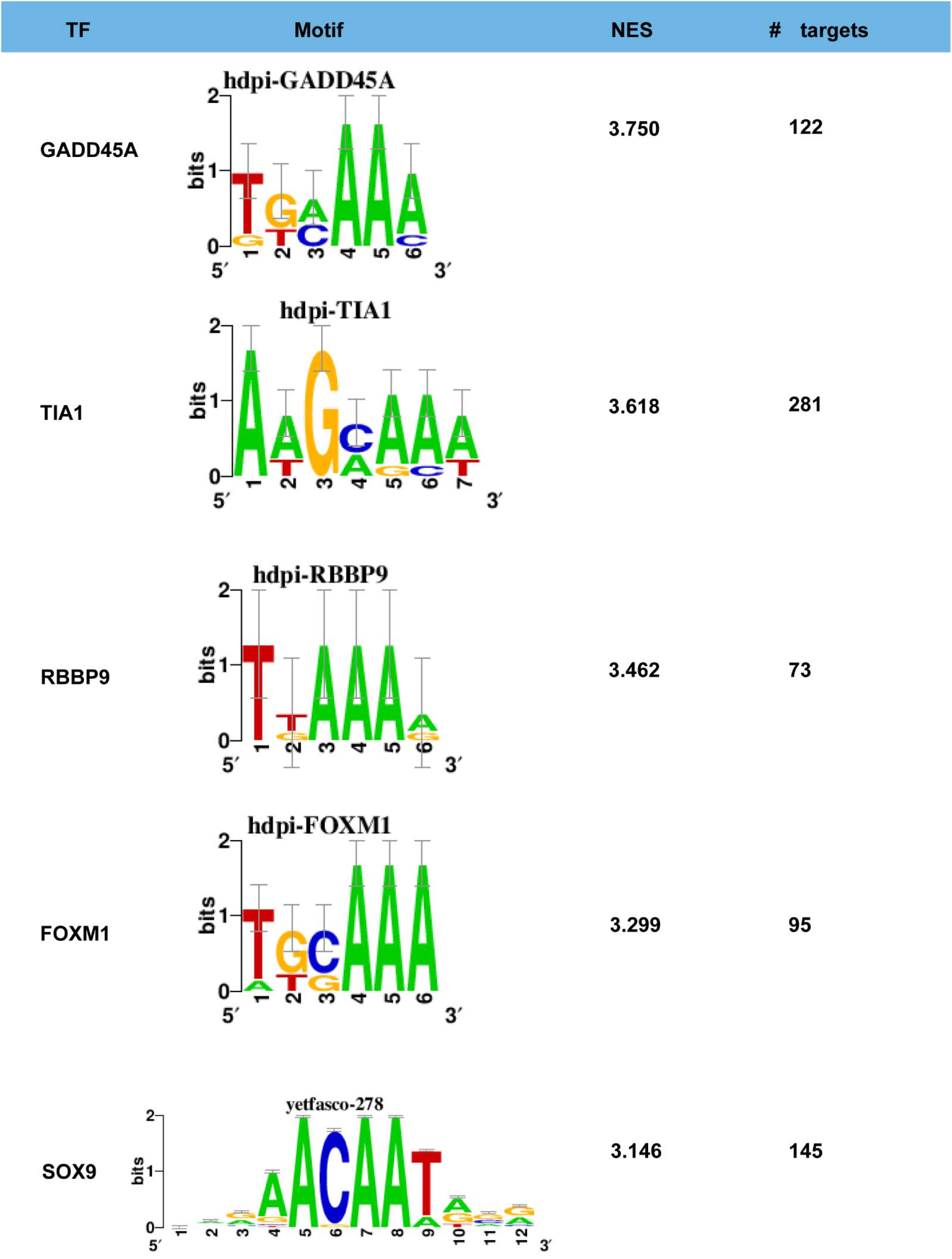
Transcription factor enrichment of upregulated genes in DMD-VSMCs compared to WT-VSMCs. Transcriptional factor, target, and motif discovery analysis showing enrichment of transcriptional factors GADD45A, TIA1, RBBP9, FOXM1 and SOX9 with iRegulon on the upregulated gene set (fold-change>2, *P<0*.*05*) in DMD-VSMCs compared to WT-VSMCs.

### Transcriptome analysis of VSMCs from normal iPSC and DMD iPSC following oxidative stress

We further analyzed the response of WT-VSMCs and DMD-VSMCs to oxidative stress. After H_2_O_2_ treatment, KEGG enrichment analysis showed upregulated genes related to Rap1 signaling pathways, AGE-RAGE signaling pathway in diabetic complications, fluid shear stress and atherosclerosis in DMD-VSMCs (**Fig.10**). The significantly enriched top 5 KEGG pathways in downregulated genes in DMD-VSMCs included ECM-receptor interaction, arrhythmogenic right ventricular cardiomyopathy, phospholipase D signaling pathway and TGF-β signaling pathway (**Fig.10**). Go enrichment of BP showed increased cell death and apoptosis, muscle contraction, reorganization of actin filament and cytoskeleton; Go enrichment of CC showed increased actin filament, sarcomeres, contractile fibers, vacuoles, vesicles, and lysosomes, endoplasmic reticulum; Go enrichment of MF showed increased binding activity to integrin, calcium, cytoskeletal protein, chemoattractant and amyloid-beta in DMD-VSMCs with oxidative stress (**Fig.11A**). While their response to hypoxia, oxidative stress, heparin, fluid shear stress was decreased; cell migration and motility were also impaired. In addition, membrane components, mitochondrion and mitochondrial protein-containing complex were downregulated. Molecular function including signaling receptor, extracellular matrix binding and transmembrane receptor protein kinase activity were impaired in DMD-VSMCs with oxidative stress (**Fig.11B**). GSEA showed negative regulation of vascular development and calcium signaling; increased aggresomes and metabolic changes in DMD-VSMCs subjected to oxidative stress (**Fig.12**).

**Fig. 10.**
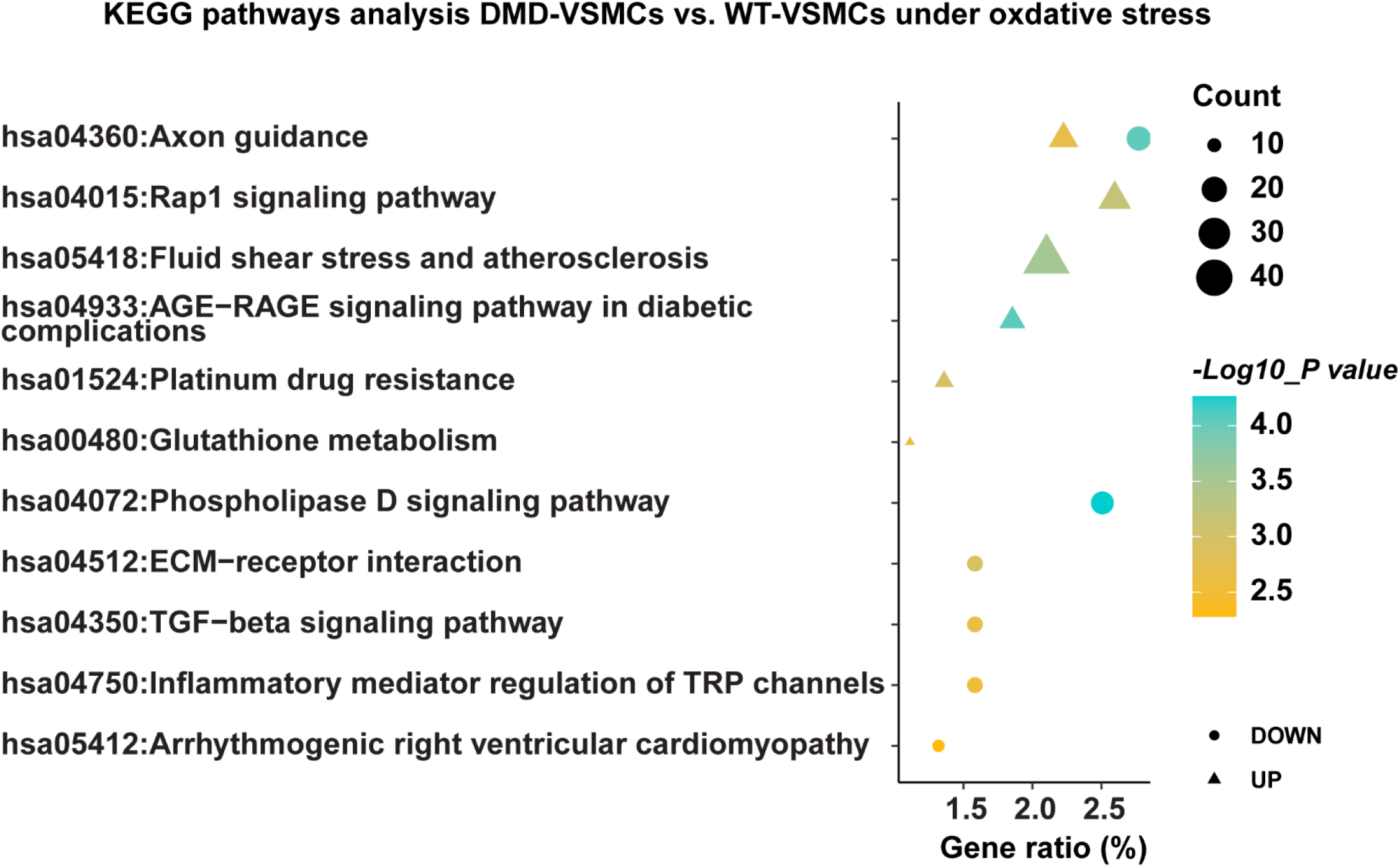
KEGG enrichment analysis for WT-VSMCs and DMD-VSMCs with oxidative stress. KEGG pathway analysis of fold enrichment from up and down regulated mRNAs with top 5 in DMD-VSMCs compared to WT-VSMCs following oxidative stress.

**Fig. 11.**
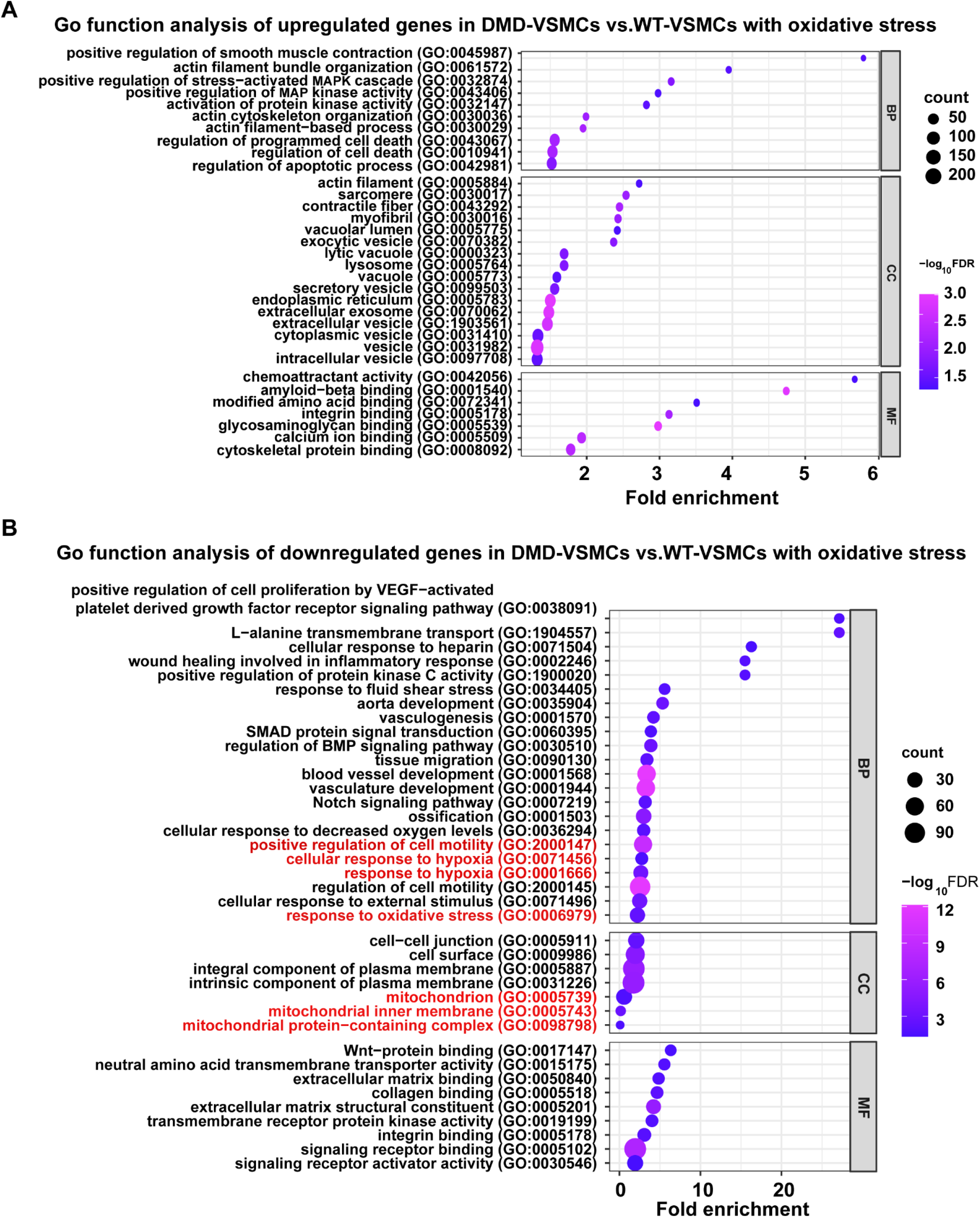
Go enrichment analysis for WT-VSMCs and DMD-VSMCs subjected to oxidative stress. Go categories of biological process (BP), molecular function (MF) and cellular components (CC) showing significant enrichment in the dataset of genes upregulated (A) and downregulated (B) in DMD-VSMCs compared to normal WT-VSMCs following oxidative stress. Low log_10_ (FDR) values are in blue and high -log10 (FDR) values are in purple, the size of the circle is proportional to the number of enriched genes.

**Fig. 12.**
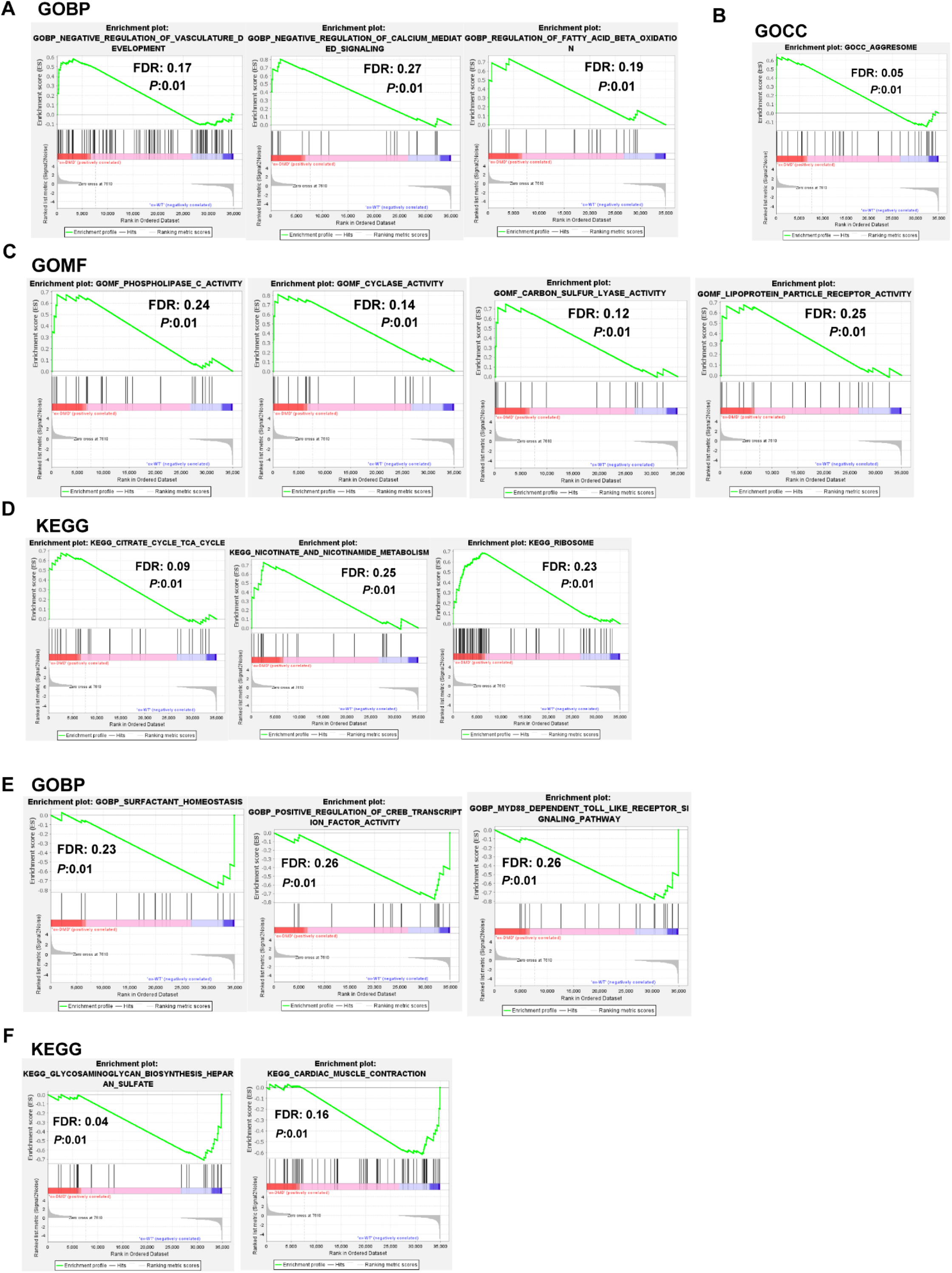
Gene set enrichment analysis (GSEA) of WT-VSMCs and DMD-VSMCs subjected to oxidative stress. GSEA biological process (BP) (A), cellular components (CC) (B), molecular function (MF) (C) and KEGG pathway (D) enrichment analysis showing significant enrichment in the dataset of genes upregulated in DMD-VSMCs with oxidative stress, and BP (E) and KEGG pathway (F) enrichment in the dataset of genes downregulated in DMD-VSMCs following oxidative stress. P<0.01, FDR≤0.25.

## Discussion

Here we report that VSMCs phenotype switching plays a critical role in multiple vascular diseases. Various stimuli can modulate the plasticity of VSMCs, such as reactive oxygen species (ROS), oxidized lipids, inflammatory cytokines, and mechanical stress ^18^. Exposure to these stimuli leads to phenotype switching of VSMCs from contraction phenotype towards synthesis phenotype, apoptosis, necrosis, degeneration, and senescence ^19^. In fact, dystrophin is expressed in normal smooth muscle cells ^5^. Tissue expression analysis showed an abundant expression of dystrophin in arteries from GTEx portal database. With single cell analysis in both mouse aorta and human aorta via single cell portal, we also showed higher expression of dystrophin in VSMCs compared with other cell types, indicating a potentially important role of dystrophin in VSMCs function. Thus, it is likely based on clinical reports that abdominal aneurysm rupture occurs in patients with Becker muscular dystrophy ^20^ perhaps due to vessel wall abnormalities. Although etiological correlation between aortic aneurysms and DMD remains unknown, this case report suggests that dystrophin deficiency may have contributed to the aortic aneurysm ^20^.

Mdx mouse is the most widely used model to study DMD, but their phenotype is milder than that of the dystrophic patients^11^. The young mdx mice show only weak DMD phenotype in skeletal muscle. A greater degree of cardiac and vessel dysfunction is observed at least in 10-12 months old mdx mice not in young mdx mice ^13^. Our study revealed that dystrophin deficiency in 12-14 months old mdx mice, resulted in spontaneous structural remodeling of the aorta. Abnormal VSMCs proliferation was accompanied by decreased expression of contractile marker Myh11, and degenerative changes supporting our view that VSMCs underwent both synthesis and degenerative phenotypes switching. These findings are in agreement with previous studies that dystrophin expression was significantly reduced during synthetic phenotype switching upon vessel injury ^14^.

Case reports on abdominal aneurysm rupture in patients with Becker muscular dystrophy suggests the involvement of major vessels in the pathogenesis of DMD disease. On the other hand, a lack of studies on VSMCs from DMD patients has been attributed to due to nonavailability of human tissue for analysis. Therefore, in this study we differentiated VSMCs from DMD iPSC to investigate the role of dystrophin deficiency in VSMCs and its mechanism. We discovered that maturation defects are present in VSMCs differentiated from DMD iPSC compared to VSMCs differentiated from normal iPSC. It is more likely that dystrophin may be a marker of VSMCs differentiation and is critical for the maintenance of contractile phenotype. It has been reported that overexpression of myocardin and myocardin-related transcription factor A (MRTF-A) promoted expression of dystrophin ^14^. However, it is obvious that dystrophin deficiency promoted loss of contractile phenotype of VSMCs. We performed the transcriptome analysis for VSMCs differentiated from human normal and DMD iPSC. KEGG enrichment analysis revealed upregulation of genes related to p53 and Hippo signaling pathways while genes related to ferroptosis, Notch signaling, osteoblast differentiation and inhibition of smooth muscle cell differentiation were downregulated. It is known that p53 signaling pathway is involved in the regulation of VSMCs proliferation ^21^. p53 activation also impaired smooth muscle differentiation via inhibition of Myocd ^22^. Hippo signaling pathway plays a critical role in the regulation of VSMCs phenotype ^23,24^. Notch signaling pathway, mediated by basic helix–loop–helix (bHLH) transcriptional repression, controls VSMCs differentiation and modulates the transcription of endogenous contractile genes in VSMCs ^25,26^. VSMCs mainly express Notch1, Notch2, and Notch3 ^27^ receptors. Jagged1-mediated Notch activation is required for the expression of smooth muscle contractility markers in VSMCs ^28-31^ In addition, mitochondria are dynamic organelles and continuously undergo fission and fusion processes. VSMCs mitochondrial metabolism has been reported as one of the mechanisms involved in the complex regulation of the VSMCs phenotype ^32,33^. GSEA showed upregulated genes related to mitochondrial depolarization in DMD-VSMCs, which was consistent with our data that disruption of mitochondrial homeostasis occurred in DMD dystrophin deficient VSMCs. Mitochondrial fission is induced by membrane depolarization ^34^. The imbalance of mitochondrial fission and fusion may contribute to the DMD-VSMCs immature phenotype. Taken together, dystrophin deficiency plays an important role in VSMC homeostasis and promoted the loss of contractile phenotype of VSMCs via dysregulation of above signaling pathways.

Oxidative stress is a prominent feature of the dystrophic pathology with increased inflammation and myonecrosis in muscle tissue ^35^. It has been reported that the lack of dystrophin renders the skeletal muscle susceptible to free radical induced injury ^36^. However, it is unclear whether dystrophin loss also affects VSMCs vulnerability to oxidative stress. Oxidative stress and oxidative DNA damage have a strong bearing on the DMD phenotype progression ^37^. In this regard, we used H_2_O_2_ treatment to recapitulate phenotype changes in DMD VSMCs in comparison with wild type VSMCs. Indeed, DMD VSMCs were highly vulnerable to H_2_O_2_ treatment with higher incidence of apoptosis. KEGG pathway enrichment analysis showed fluid shear stress and atherosclerosis increased in DMD-VSMCs. Furthermore, the biological component of the GO analysis showed the majority of enriched categories were relevant to cell death and apoptotic process, which is consistent with increased number of apoptotic cells in DMD-VSMCs exposed to oxidative stress. GO analysis also confirmed the upregulation of lytic vacuole/ vacuole and vesicle pathway in DMD-VSMCs. Per se, adult mdx mice do not exhibit the severe phenotype, but abnormalities are observed in hearts and vascular system at 10-12 months. Our results support that oxidative stress in DMD-VSMCs replicated phenotype similar to 12-14 months old mdx mice where vacuole formation propensity was increased in VSMCs suggesting that DMD iPSC derived VSMCs could be used to model disease progression in vitro by applying oxidative stress. More importantly, Go enrichment analysis also supported our idea that the response of VSMCs to oxidative stress was robust. Interestingly, cellular components including actin filament, sarcomere and contractile fibers underwent increased reorganization in DMD-VSMCs. MAPK signaling and regulation of contraction of smooth muscle cells were also upregulated by stress. However, GSEA and KEGG analysis showed that genes related to negative regulation of calcium were decreased in DMD-VSMCs affected by oxidative stress which is known to induce actin reorganization and stress fiber formation in the vascular EC and myoblasts ^38,39^. It is very likely that these filament/ fibers were stress fibers which needs further study. On a molecular scale, our GSEA analysis showed transcriptional changes in fatty acid metabolism and cholesterol homeostasis with increased genes expression related to fatty acid beta oxidation, sulfur lyase, lipoprotein particle receptor activity, nicotinate and nicotinamide metabolism in DMD-VSMCs during disease progression. However, these bioinformatic analysis derived results need further validation.

In summary, we have established in DMD iPSC derived VSMCs and mdx mouse model that the dystrophin deficiency led to VSMCs phenotype changes and disrupted mitochondrial metabolism. It is suggested that the transcriptome analysis may allow the discovery of potential signaling pathways involved in the dysregulation of transcription factors.

## Author Contributions

WX conceived the idea, participated in experimental design, acquisition, and analysis of experimental data, and drafted the manuscript. FC performed the transcriptome data analysis; XH participated in transcriptome data analysis and data assembly; SMT reviewed and edited the manuscript; MA conceived the idea, participated in experimental design, manuscript writing and editing. All authors read and approved the final manuscript.

## Data viability

The raw data of the mRNA sequence is deposited in the GEO database (GSE232219). The datasets used and/or analyzed during the current study are available from the corresponding author on request.

## Funding

Supported by NIH grant AG070145 and USF startup funds (to WX).

## Conflict of interest

None.

## Notes

### Competing Interest Statement

The authors have declared no competing interest.

